# Computational insights into mechanism of AIM4-mediated inhibition of aggregation of TDP-43 protein implicated in ALS and evidence for *in vitro* inhibition of liquid-liquid phase separation (LLPS) of TDP-43^2C^-A315T by AIM4

**DOI:** 10.1101/797613

**Authors:** Amandeep Girdhar, Vidhya Bharathi, Vikas Ramyagya Tiwari, Suman Abhishek, Usha Saraswat Mahawar, Gembali Raju, Sandeep Kumar Singh, Ganesan Prabusankar, Eerappa Rajakumara, Basant K Patel

**Affiliations:** Department of Biotechnology, Indian Institute of Technology Hyderabad, Kandi, Sangareddy, Telangana-502285, India; National Centre for Biological Sciences, Bengaluru, Karnataka, India; Virginia Commonwealth University, Richmond, USA; Department of Chemistry, Indian Institute of Technology Hyderabad, Kandi, Sangareddy, Telangana-502285, India

## Abstract

TDP-43 is an RNA/DNA-binding protein of versatile physiological functions and it is also implicated in the pathogenesis of amyotrophic lateral sclerosis (ALS) disease in addition to several other implicated proteins such as mutant SOD1 and FUS etc. Cytoplasmic mis-localization, liquid-liquid phase separation (LLPS) due to RNA depletion and aggregation of TDP-43 are suggested to be important TDP-43-toxicity causing mechanisms for the ALS manifestation. So far, therapeutic options for ALS are extremely minimal and ineffective therefore, multi-faceted approaches such as treating the oxidative stress and inhibiting the TDP-43’s aggregation are being actively pursued. In our recent study, an acridine imidazolium derivative compound, AIM4, has been identified to have anti-TDP-43 aggregation propensity however, its mechanism of inhibition is not deciphered. In this study, we have utilized computational methods to examine binding site(s) of AIM4 in the TDP-43 structure and have also compared its binding efficiency with several other relevant compounds. We find that AIM4 has a binding site in the C-terminal amyloidogenic core region of amino acids aa: 288-319, which coincides with one of the key residue motifs that could potentially mediate liquid-liquid phase separation (LLPS) of TDP-43. Importantly, alike to the previously reported effects exerted by RNA molecules, we found that AIM4 could also inhibit the *in vitro* LLPS of a recombinantly purified C-terminal fragment TDP-43^2C^ bearing an A315T familial mutation. Antagonistic effects of AIM4 towards LLPS which is believed as the precursor process to the TDP-43’s aggregation and the *in silico* prediction of a binding site of AIM4 on TDP-43 occurring in the same region, assert that AIM4 could be an important molecule for further investigations on TDP-43’s anti-aggregation effects with relevance to the ALS pathogenesis.

## Introduction

Aggregation of misfolded proteins into amyloid-like aggregates is a prominent feature of several neurodegenerative diseases such as Alzheimer’s disease, amyotrophic lateral sclerosis (ALS) and Parkinson’s disease etc. [1-5]. Amyloids can be formed by several proteins and peptides and they are ordered aggregates formed by conformational changes in the normally soluble proteins which thereafter assemble into insoluble aggregates that are partially resistant to protease degradation [1, 5-9]. The structure of amyloid aggregates consists predominantly of β-sheets arranged in a cross-β conformation where the β-strands are arranged perpendicular to the fiber axis [10]. Amyloid aggregates can bind to flat dyes such as thioflavin-T and Congo red and their binding characteristically alters the spectroscopic properties of these dyes [10-13]. Amyotrophic lateral sclerosis (ALS) is a fatal, progressive neurodegenerative disease associated with motor neuron degeneration [14-16]. Although most of the ALS cases (∼90%) are sporadic, about 5-10% of ALS cases are familial and follow Mendelian inheritance of mutations in several genes of which *SOD1, TARDBP, C9ORF72* and *FUS* are prominent [5, 17-20]. About 97% of the ALS patients harbor deposition of aggregated TAR-DNA binding protein (TDP)-43 in the disease affected tissues [5, 21]. In addition to ALS and a subset of frontotemporal lobar degeneration (FTLD-TDP-43), TDP-43 aggregates have also been found to be present in many other neurodegenerative diseases including Alzheimer’s disease and Parkinson’s disease [5, 22, 23]. TDP-43 is a ubiquitously expressed versatile RNA/DNA-binding protein and its N-terminal domain contains two tandem RRMs (RNA recognition motifs) and nuclear localization and export signals whereas its C-terminal region is a glycine-rich low complexity and intrinsically disordered domain [21, 24-26]. TDP-43 participates in a variety of cellular processes including RNA splicing, mRNA turnover, RNA trafficking, microRNA biogenesis, translation, apoptosis, neurite outgrowth and embryo development [2, 27, 28]. Also, TDP-43 undergoes caspase mediated abnormal C-terminal fragmentation which may enhance its aggregation in the ALS patients [29]. Notably, TDP-43 has been proposed to form prion-like self-seeding aggregates *in vitro* especially from its C-terminal glycine-rich region which is highly aggregation-prone [30, 31]. In fact, a fragment encompassing its RRM2 and the C-terminal region aa: 193-414 (termed: TDP-43^2C^) has been shown to have similar aggregation behavior as that of the full-length TDP-43 [30]. Also, the TDP-43^2C^ aggregates could induce aggregation of monomeric TDP-43 in cell lines *via* a prion-like seeding mechanism [30]. Recently, modulation of the *in vitro* aggregation of TDP-43^2C^ by post-translational modification and anions has been reported [32]. In addition to TDP-43^2C^, a peptide fragment, aa: 286-331, from the C-terminal region of TDP-43 has also been shown to be amyloidogenic and neurotoxic [33]. Furthermore, the aa: 311-360 segment from the C-terminal region of TDP-43 has been proposed to harbor the amyloidogenic core for inducing TDP-43 aggregation [33-35].

Mutations in TDP-43 are associated with sporadic and familial ALS [5, 18]. Notably, most of the mutations found in the ALS patients lie in the C-terminal domain, including the glycine-rich domain of TDP-43 [36]. A missense mutation A315T in the C-terminal region of TDP-43 leads to familial ALS with autosomal dominant inheritance [18, 37-39]. In fact, overexpression of the mutant human TDP-43-A315T leads to alteration in the TDP-43 expression levels, function, increased phosphorylation, fragmentation, aggregation, mitochondrial aggregation and dysfunction, ER stress, autophagy and neurodegeneration in different models [33, 40-51]. Furthermore, *in vitro* studies have revealed that a peptide fragment (aa: 286-331) having A315T familial mutation is also amyloidogenic and neurotoxic [33]. Also, in an *in vitro* study, a mutant TDP-43 A315T peptide (307-319) was found to form an anti-parallel β-sheet structure bearing thick, long and straight fibrils and was also found to be neurotoxic [52]. Recently, membrane-less liquid droplet-like organelles formed by the proteins containing prion-like low complexity domains (LCD) are being implicated in several neurodegenerative diseases including ALS [5, 53-56]. The process of formation of these membrane-less liquid droplet-like organelles is termed as liquid-liquid phase separation (LLPS) [55, 57]. Proteins containing prion-like LCDs can undergo phase-separation upon addition of salt, or alteration in pH or temperature, through transient intermolecular interactions such as the hydrophobic, cation-π and π-π interactions [58]. TDP-43 contains an LCD and can undergo LLPS in the presence of salt or due to post-translational modifications like poly ADP-ribosylation (PARylation) [59]. Strikingly, depletion of RNA concentration was found to enhance the LLPS of TDP-43 and in contrast increase in the RNA to protein stoichiometry was found to prevent and also reverse the LLPS of TDP-43 [60]. It has been proposed that LLPS is a precursor process for the formation of terminally irreversible pathogenic aggregates of TDP-43, thus, finding any molecules inhibitory to LLPS will be of high significance [61]. The aggregation of TDP-43 has been proposed to cause cytotoxicity *via* several mechanisms such as oxidative stress induction, mitotoxicity, vacuolar fragmentation and RNA metabolism dyshomeostasis etc. therefore, TDP-43 is considered as an important protein target for screening of drugs against ALS and other TDP-43 proteinopathies [5, 28, 62-64]. Thus far, no effective cure or treatment exists for ALS. A small molecule, riluzole, which is an inhibitor of glutamate mediated-excitotoxicity and another molecule edaravone (also called radicava), a potent pyrazolone free radical scavenger and an antioxidant, are the two FDA-approved drugs available thus far for ALS for selected patients and these may increase survival by approximately two to three months only [65, 66]. Current research on ALS therapeutics focuses on strategies including pro-survival signaling of motor neurons, improving mitochondrial function and anti-oxidant response and anti-aggregation approaches such as using small interfering RNA and antisense oligonucleotides [5, 67, 68]. In a recent study, towards therapeutic approach, over-expressions of the yeast co-chaperone Sis1 (Hsp40) as well as its human homolog DNAJB1 could reduce the TDP-43 toxicities respectively in the yeast cells and primary cortical neurons thereby asserting that the modulation of chaperone activity may be a potentially fruitful strategy to pursue towards ALS therapeutics [69]. In another approach, small molecules like methylene blue and dimebon were found to inhibit the TDP-43 aggregation in cell lines but these unfortunately failed further in the clinical trials [70]. In our recent study, where several imidazolium derivatives of acridine were screened, AIM4 was found to have high anti-TDP-43 aggregation effects in the yeast model expressing TDP-43-YFP and also *in vitro* as tested against a recombinantly purified C-terminal aa: 193-414 fragment (TDP-43^2C^) [30, 71, 72]. However, the mechanism by which AIM4 inhibits TDP-43 aggregation remains to be elucidated thus whether it has any binding affinity to TDP-43 needs to be determined. Also, as TDP-43 is highly aggregation-prone and needs to be maintained in presence of denaturants, it thwarts the usage of traditional tools such as isothermal titration calorimetry (ITC) to determine the AIM4’s binding affinity. Therefore, in this study computational tools like molecular docking and molecular dynamic simulations, were used to get insights into the binding mechanism of AIM4 to TDP-43. For comparison, we examined several other acridine imidazolium derivatives [71, 72] and a previously reported anti-TDP-43 aggregation molecule dimebon. Dimebon has been previously shown to exhibit anti-aggregation effects on TDP-43 in SH-SY5Y cells [70]. Another small anti-amyloidogenic molecule, DPH, that is thus far not known to influence the TDP-43’s aggregation has also been examined here for comparison [70]. As LLPS has been suggested to be a precursor process for the aggregation and deposition of TDP-43 as well as many other RNA binding proteins, therefore, we also examined whether AIM4 can inhibit the *in vitro* LLPS of the TDP-43^2C^ bearing an A315T familial ALS mutation. Together, we attempted here to investigate the potential binding site(s) of AIM4 on TDP-43 and its potential application towards inhibiting the liquid-liquid phase separation behavior of TDP-43 which is proposedly pathogenic for ALS.

## Materials and methods

### Materials

Ni-NTA agarose was purchased from Qiagen (USA). Alexa Fluor, Thioflavin-T (ThT), Tris-base, ampicillin, chloramphenicol, sodium dodecyl sulfate (SDS), sarkosyl, dithiothreitol (DTT), phenylmethanesulfonyl fluoride (PMSF), isopropyl β-D-1-thiogalactopyranoside (IPTG), and imidazole were procured from Sigma (USA). Guanidine hydrochloride (GdnHCl), TCEP ammonium persulphate (APS) were purchased from SRL (India). Bradford’s protein concentration estimation reagent was from Bio-Rad (USA). EDTA-free protease inhibitor cocktail was purchased from Roche Diagnostics (Switzerland). Agarose, Coomassie brilliant blue R-250, tetramethylethylenediamine (TEMED), sodium phosphate dibasic and sodium phosphate monobasic were purchased from HiMedia Lab (India). Urea was purchased from Affymetrix (USA). Restriction enzymes were procured from NEB BioLabs (USA). Dimebon was purchased from Sigma (USA). Diphenhydramine hydrochloride (DPH) was procured from TCI chemicals (Canada). Alexafluor 488 C5 malemide (catalogue number: A10254) was purchased from Thermofischer scientific, USA.

### Receptor and ligand preparation

#### Protein structure

In previous studies, several available structures of small segments of amyloid forming proteins have been capitalized for the design of inhibitors of fibril formation [73, 74]. In this study, we used a Cryo-EM determined structure of a small fragment of ALS familial mutant A315E TDP-43 encompassing amino acids from aa: 288-319 (PDB ID: 6N3C, originally termed as SegB) [75]. This is a fibrillar structure of a speculated pathogenic core involved in human TDP-43 aggregation which proposedly participates in the formation of both reversible and irreversible aggregates [75]. The different chains in the 6N3C fibril structure have been labeled from A to T. We introduced another familial point mutation, A315T, in the 6N3C structure, by using the Swiss PDB viewer [76] and this structure, represented hereby as 6N3C-A315T, has been used along with the structures 6N3C for docking and MDS studies. All the ligand structures were docked sequentially to the two fibrillar structures. We also used crystal structures of the potential amyloidogenic C-terminal fragments whose PDB IDs and the amino acid numbers are: 5WIA (aa: 370-375), 5WIQ (aa: 396-402), 5WKD (aa: 300-306), 6CB9 (aa: 328-333), 6CEW (aa: 321-326), 6CFH (aa: 333-343), 5WHN (aa: 312-317), 5WHP (aa: 312-317 having a point mutation A315T), 6CF4 (aa: 312-317 having A315T mutation and phosphorylated Thr-315) and 5WKB (aa: 312-317 having point mutation A315E) [25]. In addition to the C-terminal region of TDP-43, the available structures of the tandem RRM domains (PDB ID: 4BS2, aa: 96-269) [77] and N-terminal domain (NTD) (PDB ID: 2N4P, aa: 1-89) [78] were also used for the docking studies. The overall domain architecture of TDP-43, also highlighting the amyloidogenic C-terminal region peptides, has been schematically depicted in **Figure 1**.

**Figure 1:**
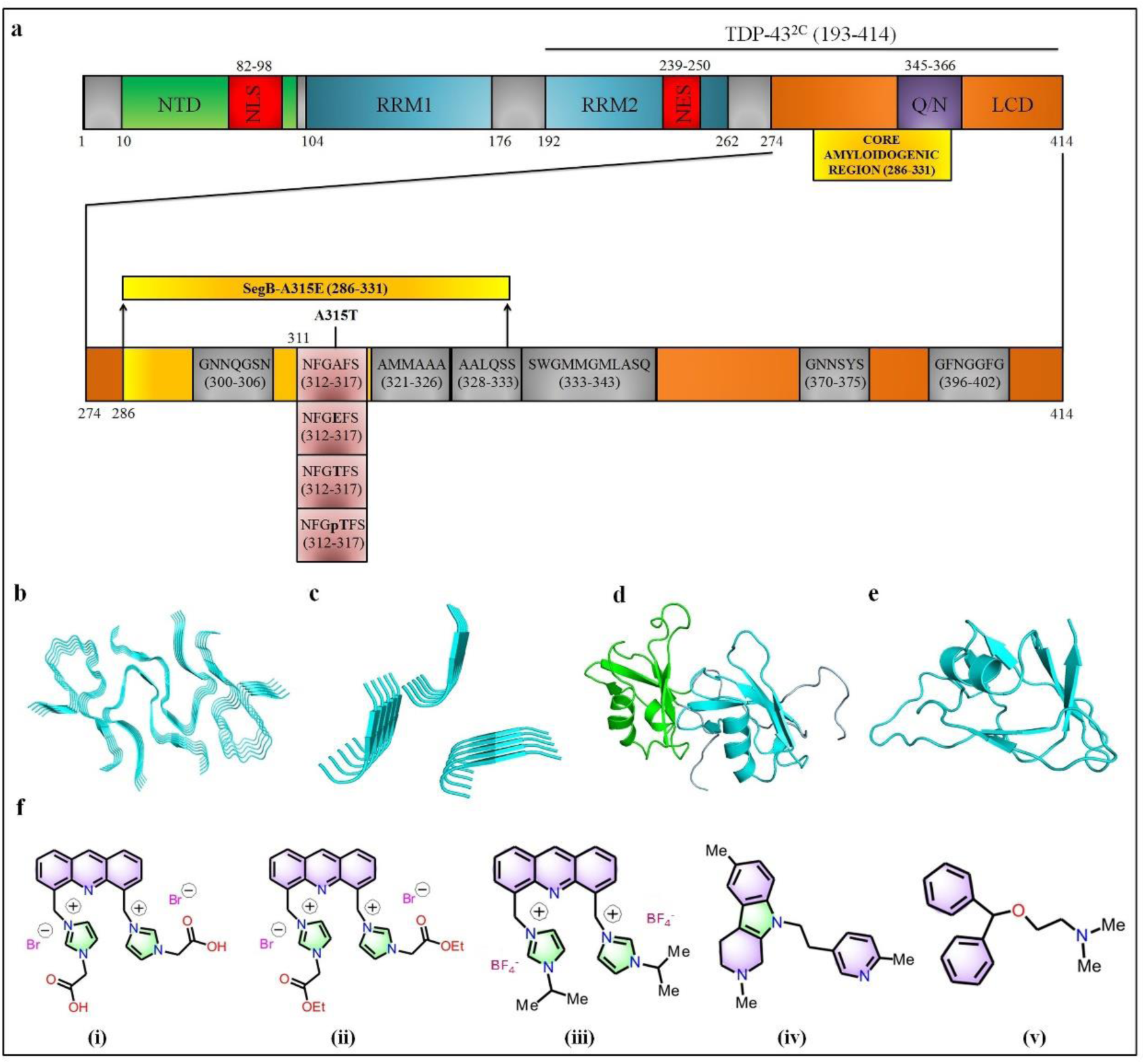
Protein and ligand structures used for docking. **a**. Schematic of the domain structure of TDP-43 which comprises of: N-terminal domain (NTD), nuclear localization signal (NLS), the two RNA Recognition Motifs (RRM1 and RRM2), nuclear export signal (NES) and a carboxy-terminal low complexity domain (LCD) with glutamine/asparagine (Q/N)-rich segment. The LCD is enlarged to depict the sequences of the ten short amyloidogenic C-terminal peptide fragments whose structures have been determined and submitted to PDB database namely 5WIA (aa: 370-375), 5WIQ (aa: 396-402), 5WKD (aa: 300-306), 6CB9 (aa: 328-333), 6CEW (aa: 321-326), 6CFH (aa: 333-343), 5WHN (aa: 312-317), 5WHP (aa: 312-317 having a point mutation A315T), 6CF4 (aa: 312-317 having A315T mutation and phosphorylated Thr-315) and 5WKB (aa: 312-317 having point mutation A315E). Sequence of the fibrillar structure of a potential core amyloidogenic region called SegB (aa: 286–331) has been depicted as yellow bar. **b**. Cryo-EM structure from the C-terminal core amyloidogenic region of TDP-43 with A315E mutation (aa: 288-319) (PDB ID: 6N3C). This structure has fibrillar morphology with multiple chains labelled from A to T. **c**. Structure of an amyloidogenic short peptide region (aa: 312-317) (PDB ID: 5WHN) **d**. Structure of the tandem RRM1 and RRM2 domains (PDB ID-4BS2) **e**. Structure of a N-terminal region (aa: 1-89) (PDB ID-2N4P) **f**. Structures of the ligands used in this study. The ligands used for the studies are- (i) AIM4: [4,5-bis{(N-carboxy methyl imidazolium)methyl}acridine] dibromide; (ii) AIM1:4,5-bisN-ethoxycarbonyl methyl imidazoliummethylacridine dibromide; (iii) AIM3: 4,5-bisN-isopropylimidazolium methylacridine tetrafluoroborate; (iv) Dimebon; (v) DPH: diphenhydramine hydrochloride.

#### Ligand structure

AIM4 and other acridine-imidazolium derivatives previously examined *in vitro* for its propensity of anti-aggregation effects on wild-type TDP-43^2C^, were used as test compounds for docking studies [71, 72]. For comparison, certain other ligands such as dimebon and diphenhydramine hydrochloride (DPH) which have already been tested in cellular models for their inhibitory activity on TDP-43 aggregation were also used [70]. Structures of the available compounds available were retrieved in sdf format from PubChem and then converted to PDB format using Open Babel [79]. For the test ligands and other ligands whose structures were not available, the structures were first drawn using Marvin Sketch (ChemAxon), and then hydrogen atoms were added and finally the structures were 3D-cleaned and saved in ‘.pdb’ format. Furthermore, all the ligand structures were geometrically optimized by using Avogadro software before docking [80]. The structures of all the ligands used for docking to TDP-43 are depicted in **Figure 1**.

#### Molecular docking

The docking studies were performed using open source software, AutoDock 4.2. ADT (AutoDock Tool), which is a graphic user interface, was used for preparing the protein and ligand structures for docking. ADT was also used to make the grid and the docking parameter files [81]. The protein and ligand structures were prepared for docking by adding gasteiger charges. Blind docking was carried out for the 6N3C, 6N3C-A315T and TDP-43 (aa: 288-319) with all the ligands. The docking space was defined by constructing a grid box. The dimensions of the grid box were set such that the entire protein structure could fit inside the grid box. Grid maps were calculated for each atom type present in the ligand. The spacing between the grid points was set to 0.669 Å. The docking parameters were set to 100 GA runs and 2,500,000 energy evaluations for each cycle. Lamarckian genetic algorithm was used for the docking [82]. For a particular conformation of the protein, the grid and docking parameters were maintained the same for all ligands. After analysis of the docking results, the protein ligand complexes were generated using ADT. Docking parameters were kept the same for the RRMs (4BS2), NTD (2N4P) and the CTD peptide structures, as described above.

#### Molecular dynamics simulation

MD simulations were performed by using Desmond package of Schrodinger (D. E. Shaw Research, New York, NY, USA) [83]. The protein-ligand complexes having the best binding energies were used for the simulations in order to check the stability and flexibility of the complexes. Before the simulation, each complex was optimized with protein preparation wizard in the Maestro 11.0 (Maestro, Schrödinger, LLC, New York, NY, 2017) by assigning proper bond orders and optimizing the hydrogen bonds. Further, the complex structures were minimized using OPLS3 force field[84]. PROPKA was used to determine the protonation status at pH 5.0 [85]. The complex structures were solvated with pre-defined single point charge (SPC) water model orthorhombic box shape (a=b=c=10Å, α=β=ϒ=90°). The entire system was neutralized by adding proper counter ions (Na^+^ or Cl^-^). The placement of ions in the solvated system was random. The salt, NaCl, was also added to the solvated system at 0.30M concentration. After building the solvated system, the system was energy minimized and relaxed using the default relaxation protocol of Desmond. This default relaxation protocol includes two rounds of energy minimization steps and four short MD simulation steps. The energy minimization was done by steepest descent method with a maximum of 2000 steps with and without restraints (force constant of 50 kcal/mol/Å on all solute atoms). The energy minimization steps followed by four short MD simulations are: 1) 12 picoseconds MD simulation at 10 K temperature in the Berendsen NVT ensemble (constant number of particles, volume, and temperature) with solute heavy atoms restrained (force constant of 50 kcal/mol/Å); 2) 12 picoseconds simulation at 10 K in the Berendsen NPT ensemble (constant number of particles, pressure, and temperature) with the same restraints on solute heavy atoms; 3) 12 picoseconds simulation in Berendsen NPT ensemble using the same restraints on solute heavy atoms in which the temperature was raised to 300 K; and 4) a final 24 picoseconds simulation in the Berendsen NPT ensemble at 300 K without restraints [86, 87]. After relaxation of the system, production MD simulation was performed using OPLS3 force field [84]. The temperature was kept at 300K using Nose-Hoover chain method [88] and pressure was kept at 1 bar using Martyna-Tobias-Klein method [89]. RESPA integrator was used for all simulations with default values : 2.0 femtoseconds time step for bonded interaction and short range non-bonded interactions whereas 6.0 femtoseconds time step for the long-range non-bonded interactions [90]. The production simulation was run in NPT ensemble for either 100 or 300 ns.

#### MD simulation analysis

The MD simulation results were analyzed using Maestro software of Schrodinger [91]. Simulation interactions diagram (SID) program available with the Desmond module was used to analyze protein-ligand RMSD, protein RMSF, ligand RMSF and protein-ligand interactions [83]. All the atoms were selected for RMSD and RMSF calculations. All the analyses were done for the entire range of the simulation time. At the end of the simulation, ligand binding energy (in kcal/mol) to its receptor was calculated using MM-GBSA tool of the Prime suite of Schrodinger (Prime, Schrödinger, LLC, New York, NY, 2017). For the energy calculation, the input partial charge of the ligand was used instead of assigning partial charges by using default force field. Protein residues within 5 Å distance of the ligand were allowed to be flexible for the energy calculation [92].

#### Plasmids and site-directed mutagenesis

Recombinant *E. coli* expression plasmid *pET15b-His-TDP-43*^*2C*^ which codes for carboxyl terminal aa: 193-414 of TDP-43, was a kind gift of Prof. Yoshiaki Furukawa, Keio University, Japan [30]. A familial ALS-linked missense point mutation (A315T) containing mutant TDP-43^2C^ expressing plasmid was generated using Q5 site-directed mutagenesis system (NEB, USA) using the *pET15b-His TDP-43*^*2C*^ plasmid as the template. The respective forward and reverse primers used for the mutagenesis were: 5’CTTTGGTACCTTCAGCATTAATCCAGCCATGATGGCTGCCGC 3’ and 5’ATGCTGAA*<u>GGTACC</u>*AAAGTTCATCCCACCACCCATATTACTAC-3’.

These primers were designed to incorporate a *KpnI* restriction enzyme recognition site (GGTACC). After *DpnI* digestion of the template DNA, the PCR product was transformed into competent *DH5α E. coli* cells (Invitrogen, USA). Then, plasmids were isolated from the obtained transformants and analyzed for positive *KpnI* digestion to ascertain for the successful mutagenesis.

#### Recombinant protein expression and affinity purification

For recombinant protein expression, the *pET15b-His-TDP-43*^*2C*^ *(A315T)* plasmid constructed here as described above, was first transformed into the expression competent *Rosetta 2 (DE3) E. coli* cells (Novagen, USA). Expressions of the mutant TDP-43^2C^-A315T protein and its affinity purification was carried out following prior published methods with minor modifications [30, 71]. Briefly, the *Rosetta 2 (DE3)* cells transformed with *pET15b-His-TDP-43-2C (A315T)* plasmid were induced for protein expression by addition of 1 mM IPTG to express the mutant TDP-43^2C^-A315T protein. Subsequently, the cells were harvested after 4 hours (O.D_600nm_=0.8) of induction and lysed by ultra-sonication in lysis buffer containing 6 M guanidine HCl (GuHCl) dissolved in phosphate buffered saline (PBS), pH 7.5 and added with 1 mM PMSF. After pre-clearing of the cell debris, the supernatant was loaded onto Ni-NTA agarose affinity chromatography column (Qiagen, Germany) pre-equilibrated with 6 M GuHCl-PBS for affinity binding of the His-tagged TDP-43^2C^-A315T. The Ni-NTA column was then washed with a buffer containing 6 M GuHCl-PBS and 10 mM imidazole to remove any non-specifically bound proteins. Next, the bound mutant TDP-43^2C^-A315T was eluted with the elution buffer containing 6 M GuHCl-PBS and 250 mM imidazole. Fractions of 500 µL each were collected and screened for the protein content by Bradford’s method and then analyzed on 10% SDS PAGE for protein homogeneity **(Supplementary Figure S3)**.

#### Amyloid-like aggregation kinetics of TDP-43^2C^-A315T and inhibition by AIM4 and AIM1

Before using for the aggregation, the 6M GuHCl in the protein sample was first replaced with 4M urea. For this, the TDP-43^2C^-A315T purified in 6M GuHCl was first diluted 10-fold in PBS pH 7.5 which causes the precipitation of TDP-43^2C^-A315T [71]. The precipitate was then pelleted by centrifugation at 14000 x g for 10 min at 4 °C and the supernatant was discarded to remove GuHCl. The pellet was then re-suspended and dissolved in 4M urea in PBS pH 7.5 and used for the aggregation following the method as described previously [71]. For examining the amyloid-like aggregation, the protein in 4M urea was first diluted to 400 µM final concentration in the aggregation buffer (PBS pH 7.5, 500 μM DTT; 2.5 M urea final) and added with 400 µM thioflavinT (ThT) and then incubated overnight with intermittent agitation in Enspire multimode microplate reader (Perkin Elmer, USA) maintained at 37 °C. The kinetic trend of the amyloid-like aggregation of mutant TDP-43^2C^-A315T was monitored by recording ThT emission fluorescence at 485 nm upon excitation at 442 nm [11, 71]. For examining their inhibitory potential towards the aggregation of mutant TDP-43^2C^-A315T, first 50 mM stock solutions of the acridine imidazolium derivative compounds AIM4 and AIM1 were prepared in 25% DMSO [71, 72]. Then the compounds were added to the aggregation solution containing 400 µM of mutant TDP-43^2C^-A315T, at a protein: AIM4/AIM1 ratio of 1:10, 1:15 or 1:20. Any inhibition of the aggregation was examined by recording the ThT emission fluorescence as described above.

#### Fluorescent labelling of mutant TDP-43^2C^-A315T for phase separation analysis

Recently, RNA-binding proteins like FUS & TDP-43 tagged with GFP have been shown to undergo liquid-liquid and liquid-solid phase separation in cellular context as well as under *in vitro* incubation when deprived of RNA molecules and this process is proposed to be relevant to the pathogenesis of ALS [60]. Thus, we examined whether AIM4 can prevent the phase separation of the recombinantly purified mutant TDP-43^2C^-A315T which contains one of the two RNA recognition motifs (i.e. RRM2) from the full-length TDP-43. For this, 1.4 mM of mutant TDP-43^2C^-A315T in 6M guanidine HCl was incubated with TCEP in the protein to TCEP ratio of 1:5 at room temperature for 30 minutes. Next, Alexafluor 488 C5 malemide was added to the protein to a final concentration of 14 mM (protein: alexafluor-1:10) and was incubated at room temperature for 2 hours. Before analysis of phase separation/aggregation, the 6M GuHCl in the protein sample was first replaced with 8M urea. For this, the TDP-43^2C^ in 6M GuHCl was first diluted 10-fold in PBS pH 7.5 which causes the precipitation of TDP-43^2C^-A315T. The precipitate was then pelleted by centrifugation at 14000 x g for 10 min at 4 °C and the supernatant was discarded to remove GuHCl. The pellet was then re-suspended and dissolved in 8M urea in PBS pH 7.5.

#### Phase separation analysis by fluorescence microscopy

For examining the phase separation, fluorescent labelled protein and unlabelled protein were mixed in the ratio of 1:50 (alexa fluor-labelled to unlabelled) and this protein mixture in 8M urea was diluted to 400 μM final concentration in the aggregation buffer (PBS pH 7.5, 500 μM DTT; 2.5 M urea final) and then incubated for 5 hours. Next, the sample was examined for phase separated condensed structures in a Leica DM2500 fluorescence microscope using the GFP filter. Images were acquired using 10x objective lens and the presence of any globular, green fluorescent positive structures, was interpreted as successful *in vitro* phase separation of the incubated soluble TDP-43^2C^-A315T proteins [60]. For examining its inhibitory potential towards phase separation of mutant TDP-43^2C^ first 50 mM stock solutions of the acridine imidazolium derivative AIM4 was prepared in 25% DMSO. Similarly, 50 mM stock concentration of DPH and dimebon were prepared in water. Then the compounds were added to the aggregation solution containing 400 µM of mutant TDP-43^2C^-A315T, at a protein: compound ratio of 1:15. Any inhibition of the phase separation was examined by fluorescence microscopy.

## Results and discussion

### Docking of AIM4 and other compounds with different known structures of TDP-43 (PDB IDs: 6N3C, 4BS2 and 2N4P)

In our previous study, we tested several acridine imidazolium derivatives and found that AIM4 was the most effective ligand to inhibit the *in vitro* aggregation of the TDP-43^2C^ [71]. AIM4 also effectively inhibited full-length TDP-43 aggregation in the yeast model. So far, the mechanism of AIM4 mediated anti-aggregation of TDP-43 is not deciphered. Thus, in an attempt to decipher the mechanism of AIM4 mediated anti-aggregation of TDP-43, we first examined the binding of AIM4 to several pre-determined structured of TDP-43, using molecular docking tool, AutoDock. Recently, the structure of a C-terminal fragment of TDP-43, which has been speculated to be a pathogenic core segment (residues 286-331 with A315E mutation) was solved using cryo-electron microscopy (PDB ID: 6N3C). To this fibrillar 6N3C structure we also performed another mutation to replace E by T to generate A315T familial ALS mutation. The two structures, 6N3C and 6N3C-A315T (288-319) were used for docking studies. In addition to the fibrillar structure 6N3C, we also used structures of TDP-43 tandem RRMs (PDB: 4BS2) and N-terminal region (PDB: 2N4P) for docking experiments.

We used the most effective anti-aggregation acridine derivative, AIM4, along with two other acridine derivatives which exhibited partial inhibitory potential towards TDP-43 aggregation, namely, AIM1 and AIM3, to carry out the docking studies with the known structures of TDP-43. As a control, we used dimebon which has been previously shown to inhibit the TDP-43’s aggregation in the cellular models and also an anti-histaminergic compound diphenhydramine hydrochloride (DPH) which did not affect the TDP-43 aggregation.

Initially, we allowed all the five ligands to bind to various TDP-43 structures by blind docking and calculated the binding energy by AutoDock. For 6N3C and 6N3C-A315T, AIM4 exhibited better binding energy over the other four ligands **(Figure 2a** and **Table 1)**. This finding that AIM4 binds better than the other ligands in the C-terminal region of TDP-43 correlates with the data obtained earlier on the *in vitro* TDP-43^2C^ structure, which also encompasses the amino acids from 288-319 of 6N3C [71].

**Table 1:**
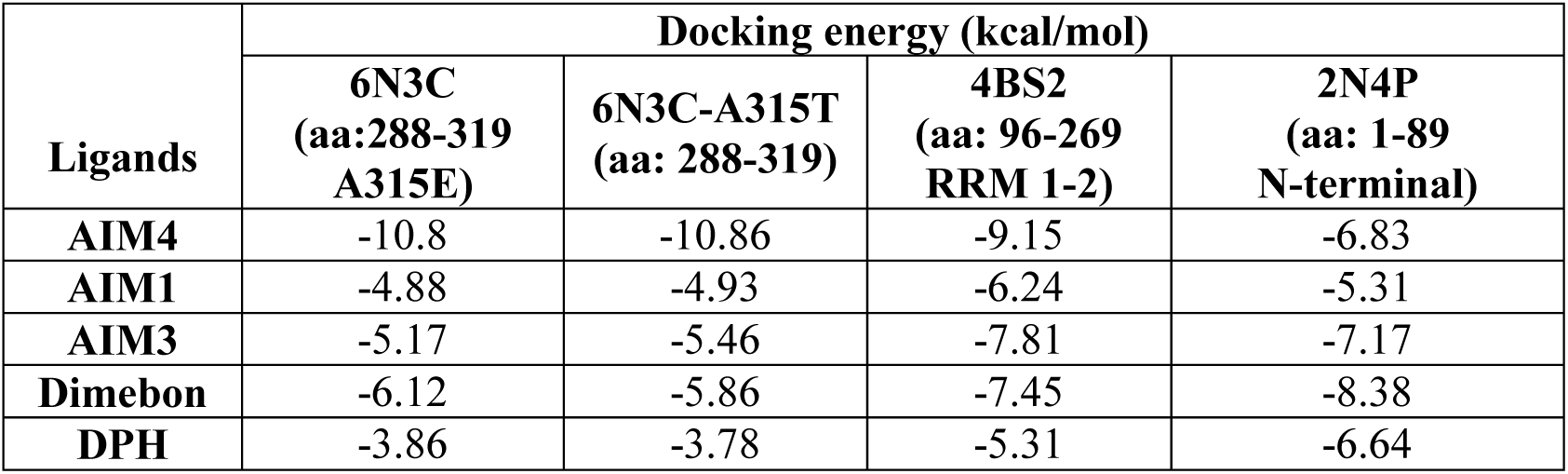
Docking energies of AIM4 and other ligands with different TDP-43 structures obtained using AutoDock.

| Ligands | Docking energy (kcal/mol) |  |  |  |
| --- | --- | --- | --- | --- |
|  | 6N3C<br>(aa:288-319<br>A315E) | 6N3C-A315T<br>(aa: 288-319) | 4BS2<br>(aa: 96-269<br>RRM 1-2) | 2N4P<br>(aa: 1-89<br>N-terminal) |
| AIM4 | -10.8 | -10.86 | -9.15 | -6.83 |
| AIM1 | -4.88 | -4.93 | -6.24 | -5.31 |
| AIM3 | -5.17 | -5.46 | -7.81 | -7.17 |
| Dimebon | -6.12 | -5.86 | -7.45 | -8.38 |
| DPH | -3.86 | -3.78 | -5.31 | -6.64 |

**Figure 2.**
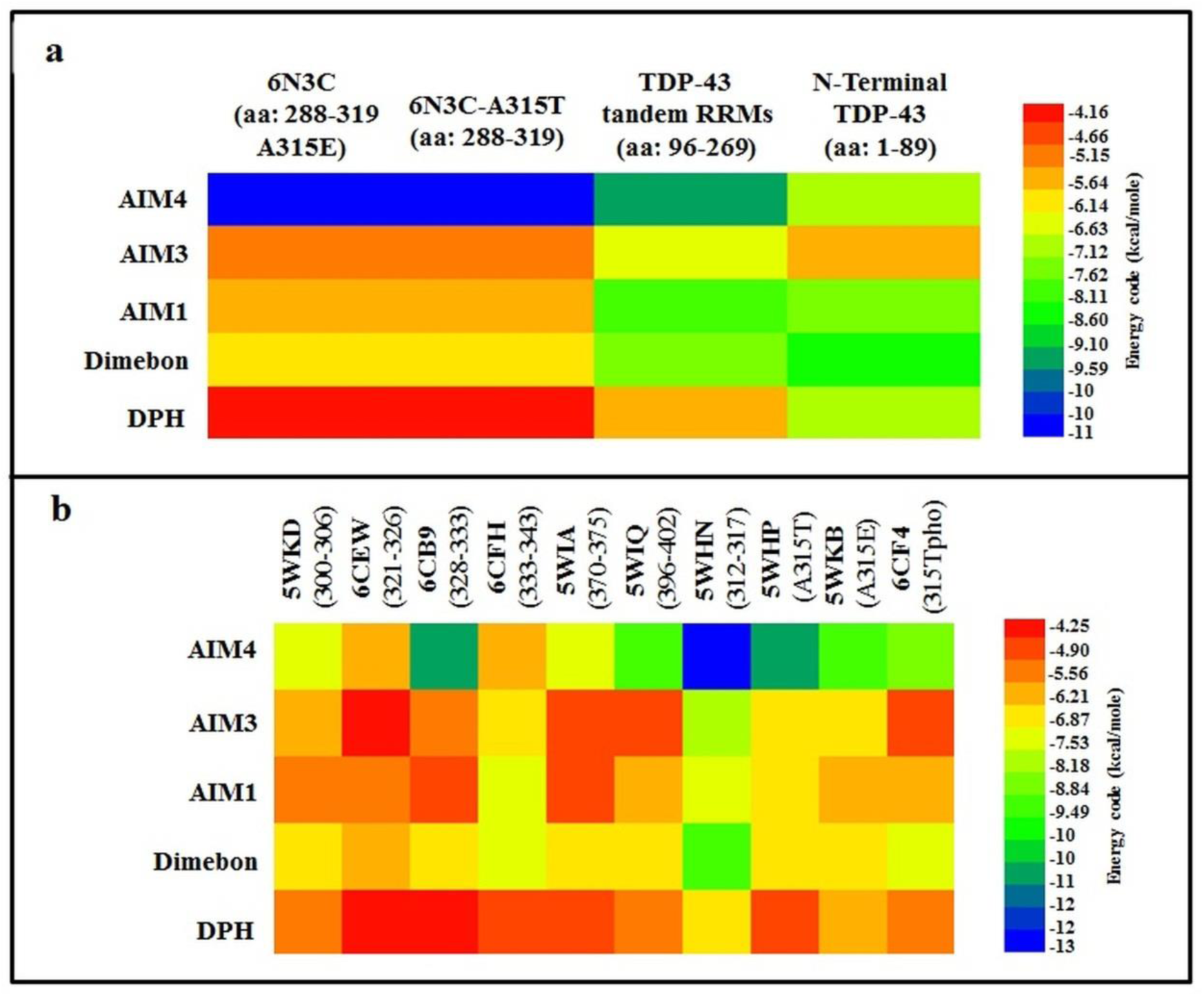
Docking energies for AIM4 and other ligands to known TDP-43 structures predicted from AutoDock. **a**. A heat map showing the docking energies of AIM4 and other ligands with A315T mutant and A315E mutant of TDP-43 (aa: 288-319) from core amyloidogenic region; TDP-43 tandem RRMs (aa: 96-269); N-Terminal TDP-43 (aa: 1-89) as obtained from rigid docking. AIM4 yielded better docking score compared to the other tested ligands with the C-Terminal region and tandem RRMs of TDP-43 structures suggesting its higher affinity. In the N-terminal region of TDP-43, dimebon and AIM3 showed better affinity that AIM4. The energy code shows regions with docking energy ranging from low (blue) to high (red). **b**. A heat map showing the docking energies of AIM4 and other ligands with short amyloidogenic peptides from the C-terminal low complexity domain (LCD) of TDP-43. Of the docking with all short peptide structures, AIM4 exhibited better binding to 5WHN peptide. The energy code shows regions with docking energy ranging from low (blue) to high (red).

Next, to examine if AIM4 has any other binding sites in the TDP-43 structure, we further performed a blind docking of all the five ligands to the structures of the TDP-43’s tandem RRMs (aa: 96-269; PDB ID: 4BS2) [77]. The tandem RRM region has high significance as TDP-43 interacts with the RNA molecules *via* the RRMs and the RNA interaction has been reported to inhibit TDP-43 from condensing into phase-separated liquid droplets, which are speculated to be precursor of its terminal irreversible amyloid-like aggregation [60]. As with the C-terminal region of TDP-43, AIM4 also exhibited better binding energy in comparison with all other ligands when docked to the tandem RRM structure **(Figure 2a** and **Table 1)**.

Additionally, the N-terminal region of TDP-43 has also been proposed to play a role in the aggregation of TDP-43 [93]. Therefore, blind docking of the ligands were also carried out against the available solution NMR structure of an N-terminal region (aa: 1-89; PDB ID: 2N4P) of TDP-43 **(Figure 2a)**. We chose the best representative conformer of 2N4P and docked it with all the ligands. In this N-terminal TDP-43 structure, dimebon yielded better docking energy than AIM4 thereby suggesting of its better binding affinity than AIM4 to the N-terminal region **(Figure 2a** and **Table 1)**. Taken together, the docking results predict that AIM4 may prefer to bind to the aggregation prone C-terminal region of TDP-43 and the tandem RRMs over the N-terminal region.

Further, it has been established that some peptide fragments from the C-terminal low complexity domain (LCD) of TDP-43 are able to *in vitro* convert into either the characteristic amyloid structures known as the steric zipper structures or into labile aggregates [25]. Hence, we examined the affinity of the ligands to these shorter TDP-43 C-terminal peptides comprising of two category of peptides: First, those capable of forming amyloid-like steric zipper structures namely, 5WKD (aa: 300-306), 6CEW (aa: 321-326), 6CB9 (aa: 328-333), 6CFH (aa: 333-343), 5WIA (aa: 370-375) and 5WIQ (aa: 396-402); and Second, those capable of forming labile aggregates, comprising of the residues 312 to 317 of TDP-43, namely, wild-type sequence (5WHN), familial mutant sequences: A315T (5WKB), A315E (5WHP), and phosphorylated A315T familial mutant (6CF4). For the peptides capable of forming amyloid-like steric zippers, 5WKD, 6CEW, 6CB9, 5WIA, and 5WIQ, the AIM4 molecule exhibited effective docking energy over the other ligands **(Figure 2b** and **Table 2)**. Similarly, for the four peptides which form labile aggregates (5WHN, 5WHP, 5WKB and 6CF4), the AIM4 molecule had better affinity than all the other tested ligands **(Figure 2b** and **Table 2)**. Among the four structures, the docking energy of AIM4 with 5WHN was predicted to be significantly better than that of the other ligands (−13.43 kcal/mol) with a predicted inhibition constant (Ki) of 0.14 nM **(Figure 2b** and **Table 2)**.

**Table 2:** Docking energies of AIM4 and the other ligands with the structure of TDP-43 short amyloidogenic peptides obtained by docking.

| Ligands | Amyloidogenic peptides from TDP-43 low complexity domain (Docking energy in kcal/mol) |  |  |  |  |  |  |  |  |  |
| --- | --- | --- | --- | --- | --- | --- | --- | --- | --- | --- |
|  | 5WKD<br>(300-306) | 6CEW<br>(321-326) | 6CB9<br>(328-333) | 6CFH<br>(333-343) | 5WIA<br>(370-375) | 5WIQ<br>(396-402) | 5WHN<br>(312-317) | 5WHP<br>(A315T) | 5WKB<br>(A315E) | 6CF4<br>(A315T<br>pho) |
| **AIM4** | -7.25 | -6.53 | -10.34 | -6.43 | -7.7 | -9.2 | -13.43 | -10.78 | -9.65 | -9.01 |
| **AIM1** | -6.08 | -4.59 | -5.46 | -6.89 | -4.97 | -5.05 | -7.96 | -6.73 | -6.86 | -4.97 |
| **AIM3** | -5.6 | -5.47 | -5.15 | -7.66 | -5.16 | -6.39 | -7.71 | -6.82 | -6.13 | -6.16 |
| **Dimebon** | -6.98 | -6.32 | -6.59 | -7.38 | -7.09 | -6.6 | -9.31 | -7.16 | -6.79 | -7.45 |
| **DPH** | -5.56 | -4.07 | -4.1 | -4.75 | -5.29 | -5.56 | -6.66 | -5.23 | -6.11 | -5.71 |

### MD simulation suggests that AIM4 forms stable complex with 6N3C and 6N3C-A315T TDP-43 (aa: 288-319)

Next, we analyzed the conformational changes of the AIM4 bound TDP-43 structure complexes through molecular dynamic (MD) simulations. First, we subjected the complex of AIM4 with 6N3C (TDP-43 aa: 288-319 A315E mutant) to molecular dynamics simulation for a period of 300 nanoseconds **(Figure 3)**. The MD simulation revealed that the trend of the mutant 6N3C-AIM4 (protein-ligand) RMSD against the simulation time was found to plateau and the average protein-ligand RMSD (2.8±0.8 Å) was nearly same as that of protein alone RMSD (2.6±0.2 Å), thereby suggesting that the complex is stable **(Figure 3a)**. Along with RMSD, protein-RMSF (root mean square fluctuation) and ligand-RMSF were also monitored to assess the local residue flexibility of the protein and the atom-wise fluctuations in the ligand **(Figure 3b)**. Total number of contacts that were made between the mutant A315E TDP-43 peptide and AIM4 and maintained throughout the course of simulation time are depicted **(Figure 3c)**. Gly-288 and Asn-291 were found to be the key residues interacting with AIM4 throughout the simulation period and these interactions were mediated majorly through H-bond **(Figure 3d)**. Over the time course of MD simulation, the protein-ligand contacts retained throughout the simulation period, have been shown as a timeline representation of number of contacts made by each residue with the ligand in each trajectory frame **(Figure 3e)**.

**Figure 3:**
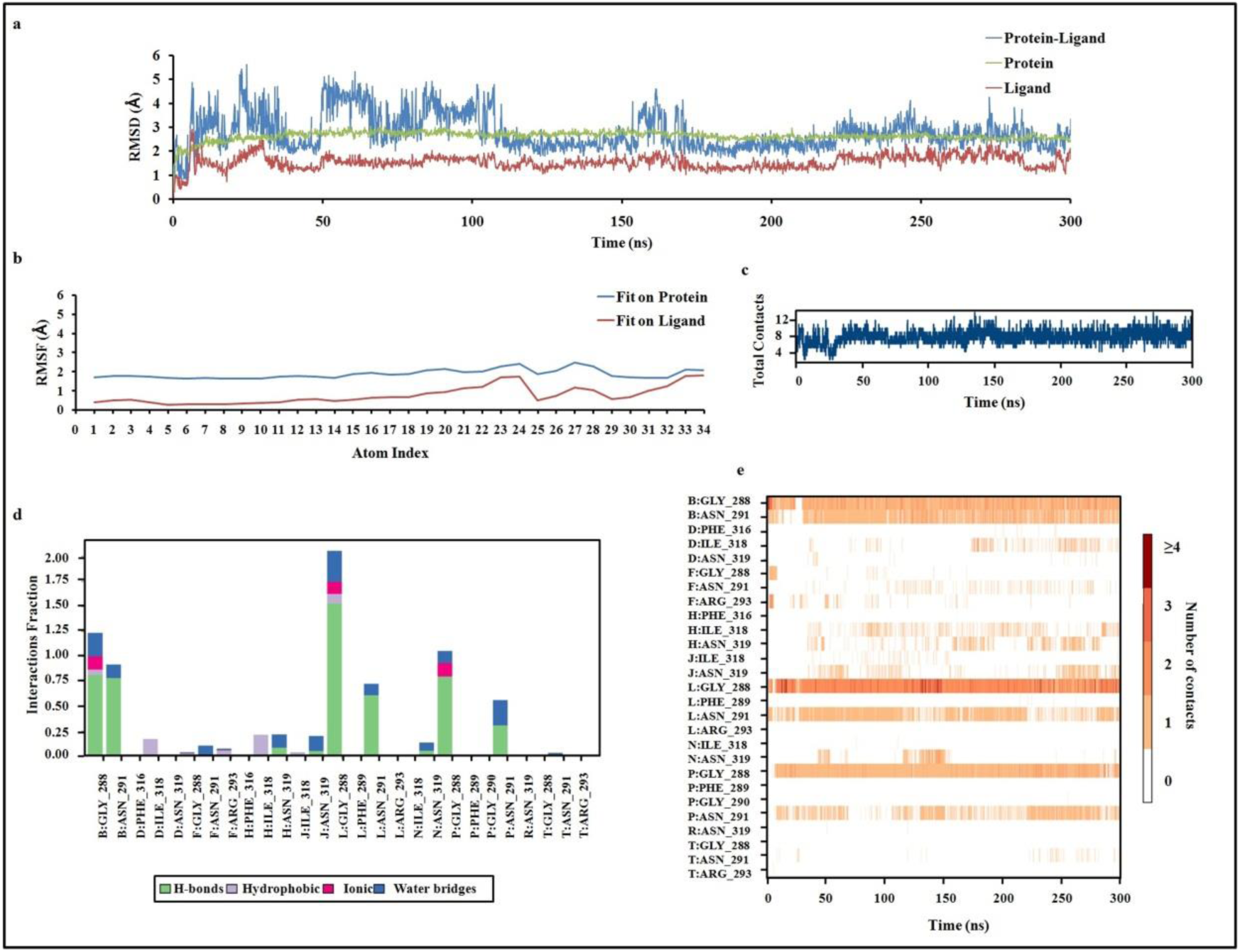
Conformational stability of 6N3C (TDP-43 aa:288-319 A315E mutant) complexed with AIM4 as determined by 300 ns of MD simulations. **a**. Protein only RMSD, protein-ligand (AIM4) RMSD and ligand only RMSD of the mutant A315E TDP-43 structure. RMSD of protein-ligand during MD simulation indicates the RMSD of a ligand when the protein-ligand complex is first aligned on the protein backbone of the reference and then the RMSD of the ligand heavy atoms is measured. Ligand RMSD indicates the RMSD of the ligand that is aligned and measured on its reference conformation. Protein only RMSD indicates the RMSD of the protein alone (Cα), where protein frames are first aligned on the reference frame backbone, and then the RMSD is calculated based on the atom selection. **b**. Root Mean Square Fluctuations (RMSF). ‘Fit Ligand on Protein’ trend indicates the ligand fluctuations with respect to the protein. ‘Ligand’ trend shows the fluctuations where the ligand in each frame is aligned on the ligand in the first reference frame **c**. The number of total contacts (H-bonds, Hydrophobic, Ionic, Water bridges) made between the mutant A315E TDP-43 peptide and the ligand (AIM4) over the course of the simulation. **d**. The interactions of AIM4 with the amino acids of mutant A315E TDP-43 peptide (aa: 288-319; PDB ID: 6N3C). The stacked bar charts have been normalized over the course of the trajectory: for example, a value of 1.0 suggests that the specific interaction is maintained for 100% of the simulation time. **e**. A timeline representation of the interactions and contacts (H-bonds, Hydrophobic, Ionic, Water bridges) showing the residues of mutant A315E TDP-43 interacting with the ligand (AIM4) in each trajectory frame. Some residues make more than one specific contact with the ligand, which is represented by a darker shade of orange, according to the scale to the right of the plot. Fibrillar structure of 6N3C contains several chains from A to T which are appropriately labeled for its AIM4 interaction.

Next, we analyzed the MD simulation of the complex of mutant 6N3C-A315T with AIM4 to assess the stability of the complex. The RMSD of the complex was found to be stabilized after around 80^th^ nanosecond (**Figure 4a**). The trend of the 6N3C-A315T-AIM4 (protein-ligand) RMSD against simulation time showed that the complex is stable (**Figure 4a**). The average protein-ligand RMSD (2.9±0.8 Å) was nearly same as that of protein alone RMSD (2.6±0.2 Å). Protein-RMSF (root mean square fluctuation) and ligand-RMSF were also monitored to assess the local residue flexibility of the protein and the atom-wise fluctuations in the ligand **(Figure 4b)**. Total contacts which were maintained throughout the simulation time have also been depicted **(Figure 4c)**. Gly-288 and Phe-289 were found to be the key residues interacting with AIM4 throughout the simulation period **(Figure 4d)**. These interactions were mediated majorly through hydrogen and hydrophobic bonds. Over the time course of MD simulation, the protein-ligand contact was retained throughout the simulation period as shown by timeline representation of number of contacts made by each residue with the ligand in each trajectory frame **(Figure 4e)**. Notably, key residues for LLPS contain a GFG motif and from the results of the simulation it is seen that glycine-288 and phenylalanine-289 are the key residues through which AIM4 interacts.

**Figure 4:**
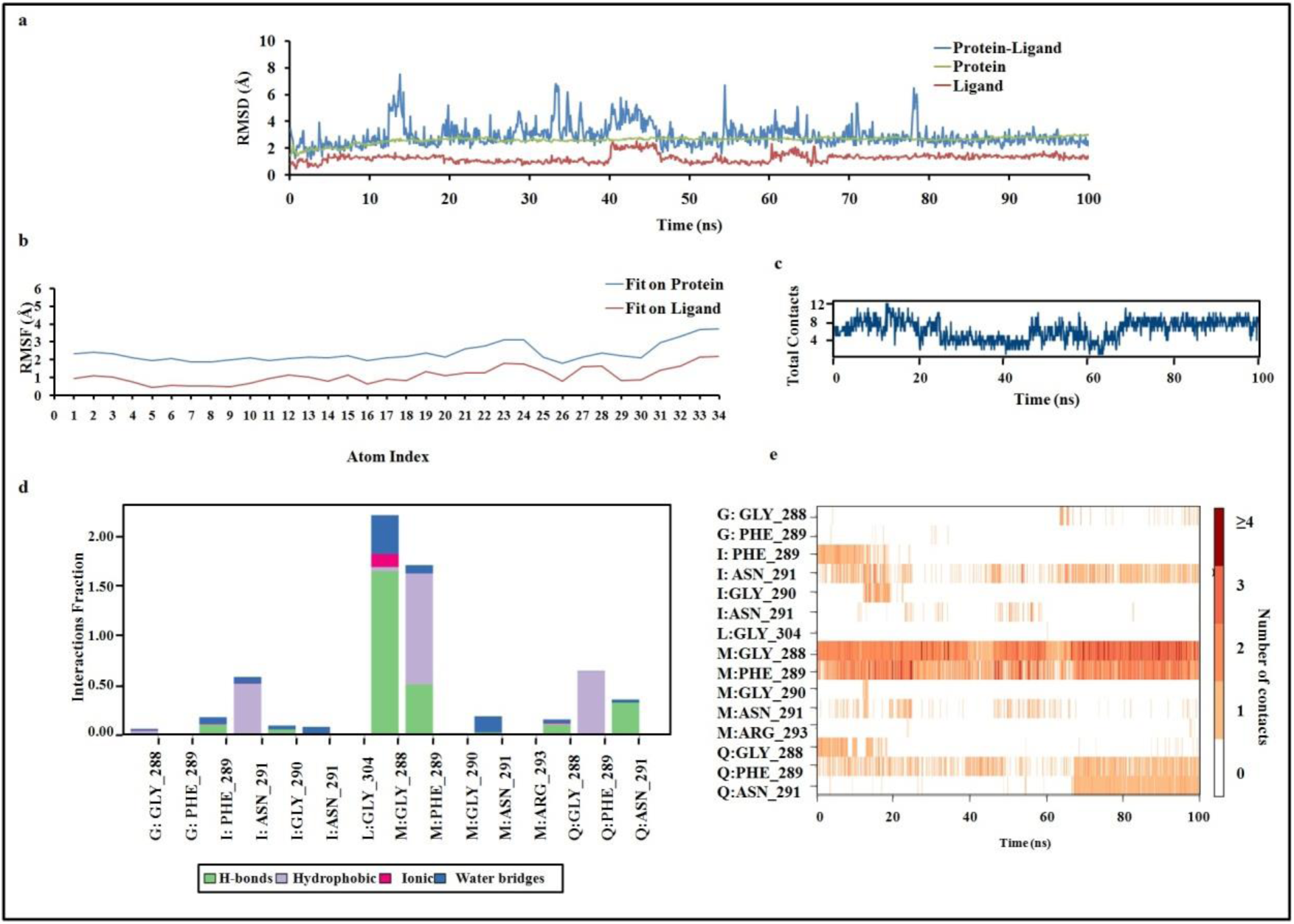
Conformational stability of the mutant 6N3C-A315T TDP-43 (288-319) complexed with AIM4 as determined by 100 ns of MD simulations. **a**. Protein only RMSD, protein-ligand (AIM4) RMSD and ligand only RMSD of the mutant A315T TDP-43 structure. RMSD of protein-ligand during MD simulation indicates the RMSD of a ligand when the protein-ligand complex is first aligned on the protein backbone of the reference and then the RMSD of the ligand heavy atoms is measured. Ligand RMSD indicates the RMSD of the ligand that is aligned and measured on its reference conformation. Protein only RMSD indicates the RMSD of the protein alone (Cα), where protein frames are first aligned on the reference frame backbone, and then the RMSD is calculated based on the atom selection. **b**. Root Mean Square Fluctuations (RMSF). ‘Fit Ligand on Protein’ trend indicates the ligand fluctuations with respect to the protein. ‘Ligand’ trend shows the fluctuations where the ligand in each frame is aligned on the ligand in the first reference frame **c**. The number of total contacts (H-bonds, Hydrophobic, Ionic, Water bridges) made between the mutant A315T TDP-43 peptide and the ligand (AIM4) over the course of the simulation. **d**. The interactions of AIM4 with the amino acids of mutant A315T TDP-43 peptide (aa: 288-319; PDB ID: 6N3C). The stacked bar charts have been normalized over the course of the trajectory: for example, a value of 1.0 suggests that the specific interaction is maintained for 100% of the simulation time. **e**. A timeline representation of the interactions and contacts (H-bonds, Hydrophobic, Ionic, Water bridges). it shows the residues of mutant A315T TDP-43 interacting with the ligand (AIM4) in each trajectory frame. Some residues make more than one specific contact with the ligand, which is represented by a darker shade of orange, according to the scale to the right of the plot. Fibrillar structure of 6N3C contains several chains from A to T which are appropriately labeled for its AIM4 interaction.

Next, to assess the stability of the AIM4 binding to the RRMs of TDP-43 (aa: 96-269; PDB ID: 4BS2) complex, MD simulation of 4BS2-AIM4 complex was carried out for 100 nanoseconds. The RMSD plot of the protein-ligand complex did not plateau and displayed increasing trend towards the end of simulation indicating that the complex was not stable. Over the 100 ns of simulation period, the average RMSD values for protein alone (6.7±2.5 Å) was observed to be very high indicating that the protein is undergoing a large conformational change during the simulation. Also, the high average value of protein-ligand (8.2±2.9 Å) RMSD suggested that the 4BS2-AIM4 complex is unstable **(Supplementary Figure S1)**.

Next, we checked the stability of the 5WHN-AIM4 peptide-ligand complex, the peptide for which AIM4 displayed maximum affinity in docking, by running MD simulations for 100 nanoseconds. For the 5WHN-AIM4 complex, high fluctuations were observed in the protein only and protein-ligand RMSD values indicating that the complex is not stable. The average protein alone RMSD and the protein-ligand RMSD were observed to be 10.1±3.5 Å and 8.7±3.7 Å, respectively **(Supplementary Figure S2)**. Thus, MD simulation studies suggest of more stable binding of AIM4 with the mutant A315E and A315T TDP-43 aa: 288-319 structures in comparison to the TDP-43’s tandem RRMs and the 5WHN peptide. In summary, the results from docking and MD simulation of 6N3C-AIM4, 6N3C-A315T-AIM4, 5WHN-AIM4 and 4BS2-AIM4 suggested one binding site for AIM4 on TDP-43 in the 288-319 region. This may be a contributing reason for the previously observed [71] better *in vitro* inhibitory capability of AIM4 on the aggregation of TDP-43^2C^.

### AIM4 binding poses and the predicted inhibitor constants for AIM4 and the other ligands with 6N3C and 6N3C-A315T TDP-43

From the MD simulation results, the binding poses and interactions of the best efficient ligand AIM4 have been shown with the 6N3C (Frame: 300 ns) and the mutant 6N3C-A315T TDP-43 (aa: 288-319) (Frame: 100 ns) **(Figure 5)**. Molecular interactions of AIM4 with 6N3C and the mutant 6N3C-A315T TDP-43 (aa: 288-319) are slightly different. In case of 6N3C, the carboxylic group of AIM4 interacts with main chain of Gly-288 and side chain of Asn-291 through hydrogen bond whereas in 6N3C-A315T mutant TDP-43 (aa: 288-319), interactions with Gly-288 and Asn-291 remains as such and also the imidazole ring of AIM4 formed pi-stacking interactions with the side chain of Phe-289.

**Figure 5:**
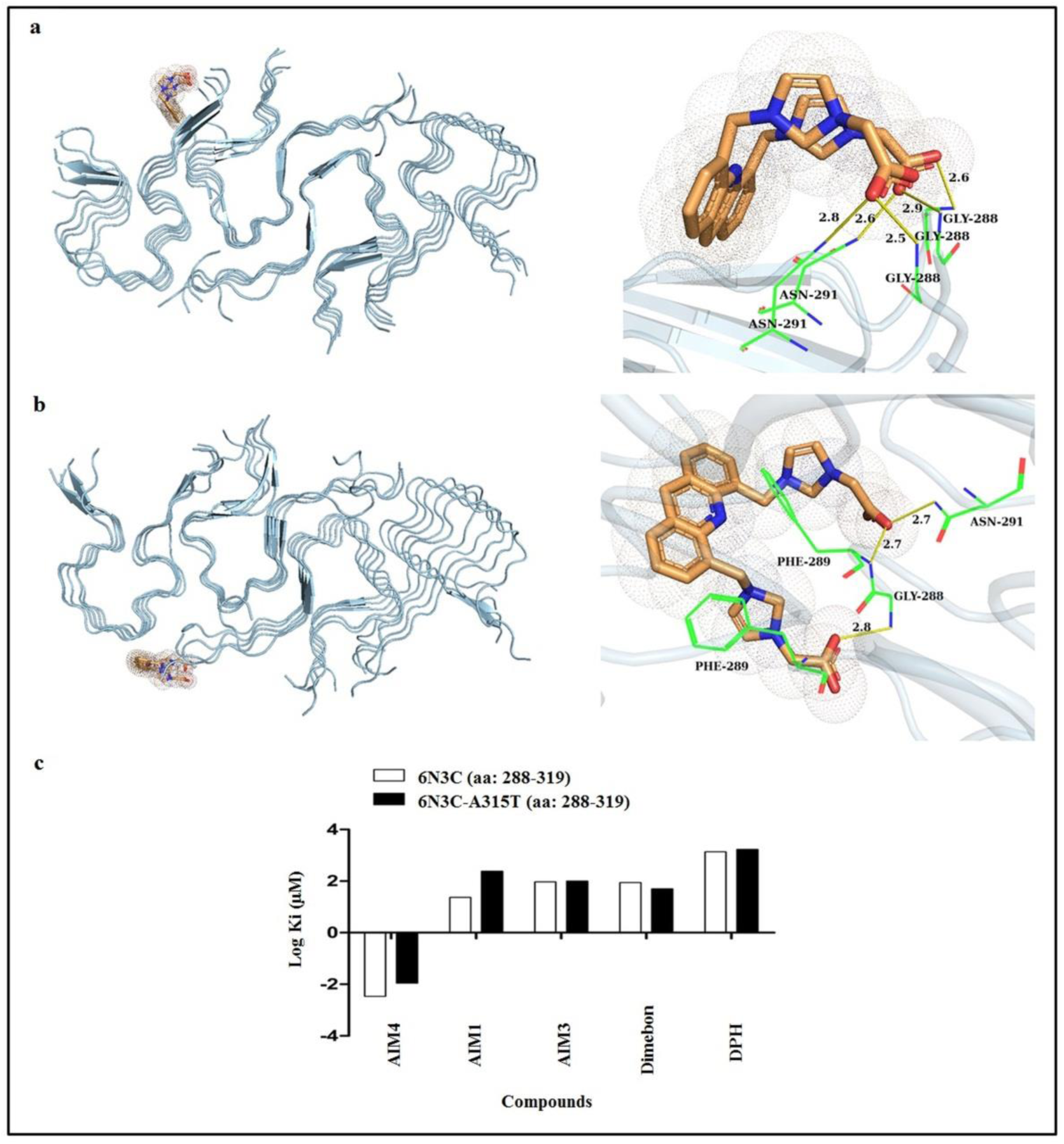
Binding poses of AIM4 and the predicted inhibitor constants for AIM4 and other used ligands with 6N3C and 6N3C-A315T TDP-43. **a**. The left panel shows AIM4 (orange) docked to 6N3C TDP-43 (aa: 288-319 A315E mutant) (grey) and the right panel displays the residues involved in molecular interactions with AIM4. **b**. The left panel shows AIM4 (orange) docked to mutant 6N3C-A315T TDP-43 (aa: 288-319) (grey) and the right panel displays the residues involved in molecular interactions with AIM4. **c**. The predicted inhibitor constants for different ligands as determined by AutoDock software.

### Binding free-energy calculations of 6N3C-AIM4 and 6N3C-A315T-AIM4 complexes from MD simulations

The binding free-energies of the complexes of the ligands with 6N3C or 6N3C-A315T mutant TDP-43 (288-319) were averaged at the end of the simulations over the last 25 ns using MMGBSA approach (Prime package of Schrodinger) [94]. The average binding free energy for the AIM4 with 6N3C was found to be −52.38 kcal/mol and in case of AIM4 with 6N3C-A315T was −52.78 kcal/mol.

### AIM4 also inhibits amyloid-like aggregation of TDP-43^2C^-A315T

Since, AIM4 binding was stable with the familial mutant A315T and exhibited better docking energy, we decided to investigate the effects of AIM4 on an ALS-related familial mutant of TDP-43, A315T mutant of TDP-43^2C^ was generated and examined for its *in vitro* aggregation kinetics. As a control, we also tested the second best acridine TDP-43^2C^ inhibitor AIM1 for comparison. Similar to as reported for the wild-type protein [71], TDP-43^2C^-A315T also showed ThT fluorescence kinetics akin to that of an amyloid-like aggregation (**Figure 6**). Furthermore, alike the wild-type TDP-43^2C^ protein [71], concentration-dependent inhibition of the aggregation of TDP-43^2C^-A315T was also observed in the presence of AIM4 and AIM1 (**Figure 6**).

**Figure 6:**
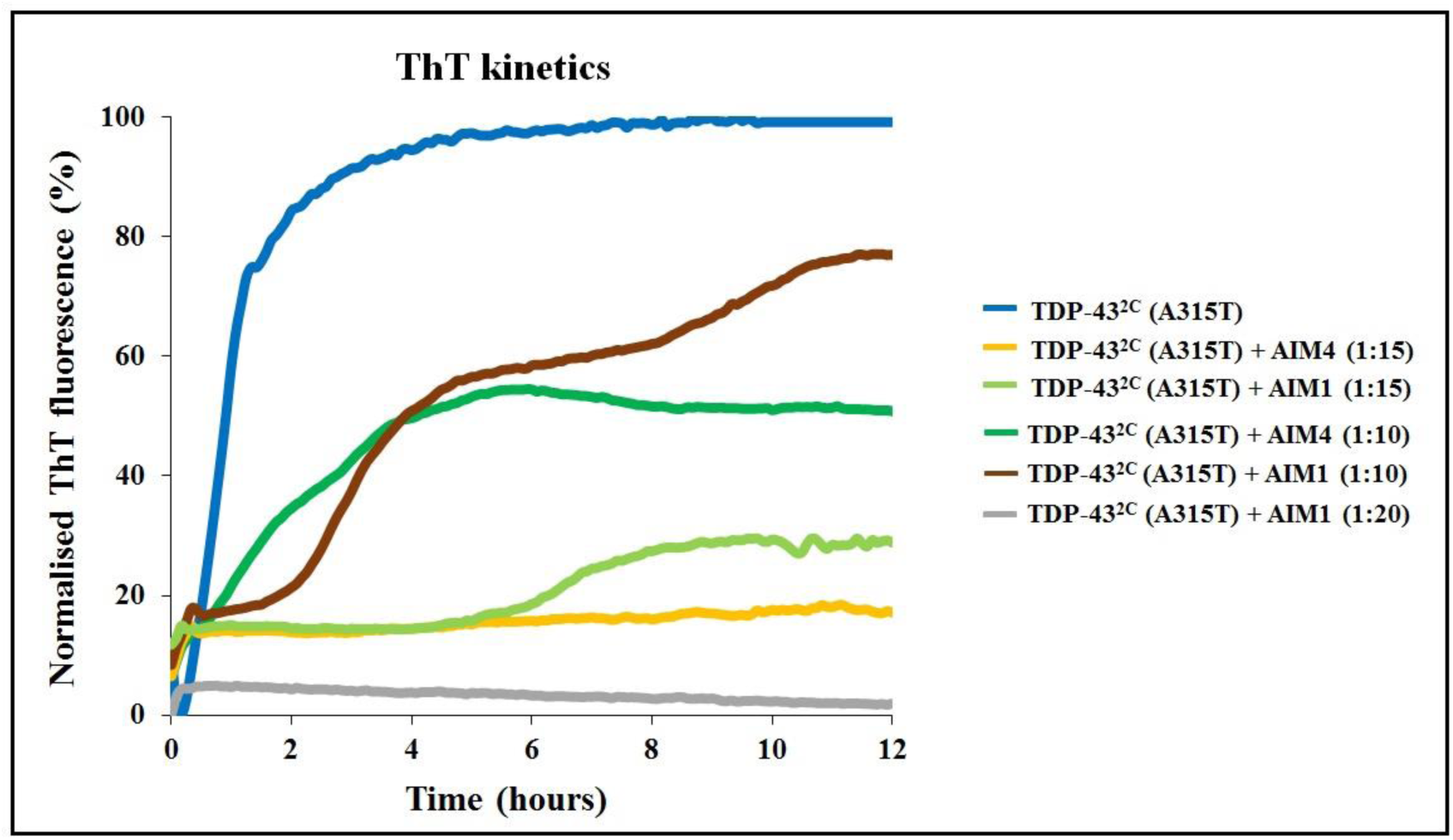
AIM4 inhibits amyloid-like aggregation of TDP-43^2C^-A315T. Inhibitions of the aggregation kinetics of the mutant TDP-43^2C^-A315T by AIM4 & AIM1 were monitored by ThT fluorescence emission. Mutant TDP-43^2C^-A315T (400 μM) was incubated at 37 °C in aggregation buffer (2.5M urea-PBS), either alone or with the additions of 1:10, 1:15 or 1:20 of AIM4 or AIM1 (protein: compound molar ratio) and the ThT fluorescence emission intensity was recorded for 12 hours at 485 nm upon excitation at 445 nm. The additions of AIM4 and AIM1 were observed to decrease the TDP-43^2C^-A315T’s aggregation in concentration-dependent manners.

The kinetic trend was observed to flatten to baseline at 1:15 protein: AIM4 concentration, whereas at 1:15 protein: AIM1 concentration the mutant TDP-43^2C^ continued to exhibit amyloid-like kinetics behavior although with a relatively prolonged lag phase as compared to the protein sample with 1:10 ratio of AIM1. However, at its further increased concentration (1:20), AIM1 also caused complete inhibition of the mutant TDP-43^2C^**-**A315T as observed by the near base line ThT fluorescence trend (**Figure 6**).

### AIM4 can also inhibit the *in vitro* liquid-liquid phase separation (LLPS) of the TDP-43^2C^-A315T protein

Liquid-liquid phase separation (LLPS) of TDP-43 is influenced by both hydrophilic and hydrophobic residues. The (G/S)-(F/Y)-(G/S) motifs promote the phase separation through transient interactions in several intrinsically disordered proteins like TDP-43 [5, 95]. From our *in silico* studies, in the 6N3C-A315T mutant TDP-43 structure (aa: 288-319), AIM4 was predicted to interact with the major amino acids, glycine and phenylalanine, which are proposedly involved in its pathogenic LLPS [95]. Therefore, the effect of AIM4 on the *in vitro* LLPS of mutant TDP-43^2C^-A315T, that also encompasses the aa: 288-319, was checked. Several reports have previously shown that TDP-43 and its prion-like C-terminal domain can undergo LLPS which can be modulated by the presence of salt, RNA or post-translational modifications like the addition of poly(ADP-ribose) [59, 60, 95, 96]. Recently, alexafluor labelled TDP-43 has been shown to form phase separated droplets [97]. In agreement, we found that after incubation under aggregation conditions, alexafluor labelled mutant TDP-43^2C^-A315T underwent transition from a soluble state to form liquid-droplets, when observed under fluorescence microscopy **(Figure 7)**. To investigate if the acridine compounds AIM4, AIM1 or AIM3 could also modulate the liquid-liquid phase separation behavior of TDP-43^2C^, *in vitro* phase separation assay was performed in the presence of these acridine compounds (protein: compound molar ratio-1:15). We observed that indeed the presence of AIM4 inhibited the formation of liquid droplet assemblies (i.e LLPS) in the mutant TDP-43^2C^-A315T whereas, the samples lacking AIM4 manifested numerous liquid droplets as previously reported by other groups **(Figure 7a)**. When the presence of liquid phase-separated droplets was examined, there was no visible phase separation in the TDP-43^2C^-A315T sample incubated along with AIM4 at 1:15 ratio of the protein to AIM4. In comparison AIM1 could greatly reduce the number of phase separated liquid droplets but unlike AIM4, AIM1 could not completely abrogate LLPS (**Figure 7a** and **7b**). Furthermore, AIM3 could not inhibit the droplet formation of TDP-43^2C^-A315T and instead several irregular shaped solid aggregates were observed in presence of AIM3. Similarly, incubation of mutant TDP-43^2C^ with DPH and dimebon also did not inhibit phase separation and rather resulted in the formation of irregular shaped aggregates possibly suggesting Liquid-Solid phase separation. This observation that AIM4 and not the other tested acridine compounds, which showed the maximum inhibitory potential against TDP-43^2C^-A315T aggregation also lead to the decrease in the phase separation of TDP-43^2C^-A315T, corroborates the specificity of AIM4 binding to TDP-43. This also bears a parallel to the ability of RNA molecules in regulating the phase separation behavior of RNA binding prion-like proteins such as TDP-43 and FUS in a concentration-dependent manner possibly by keeping the bound proteins in soluble state and thereby preventing their phase separation and their consequent aggregation [60]. In fact, injection of RNA molecules into cells has been reported to revert the protein droplet clusters in the nucleus back into the soluble state. Earlier, it has been reported that the A315T mutation increases the cytoplasmic mis-localization, aggregation and toxicity of TDP-43 in neurons [98]. Taken together, our results suggest that AIM4 binds to one of the potential LLPS motifs of TDP-43 i.e the GFG motif in the amyloidogenic region and in subsequently AIM4 helps in reducing the LLPS of TDP-43^2C^-A315T *in vitro*. Further investigations on AIM4 could potentially find its usage as anti-LLPS agent towards RNA binding protein with relevance to ALS therapeutics.

**Figure 7:**
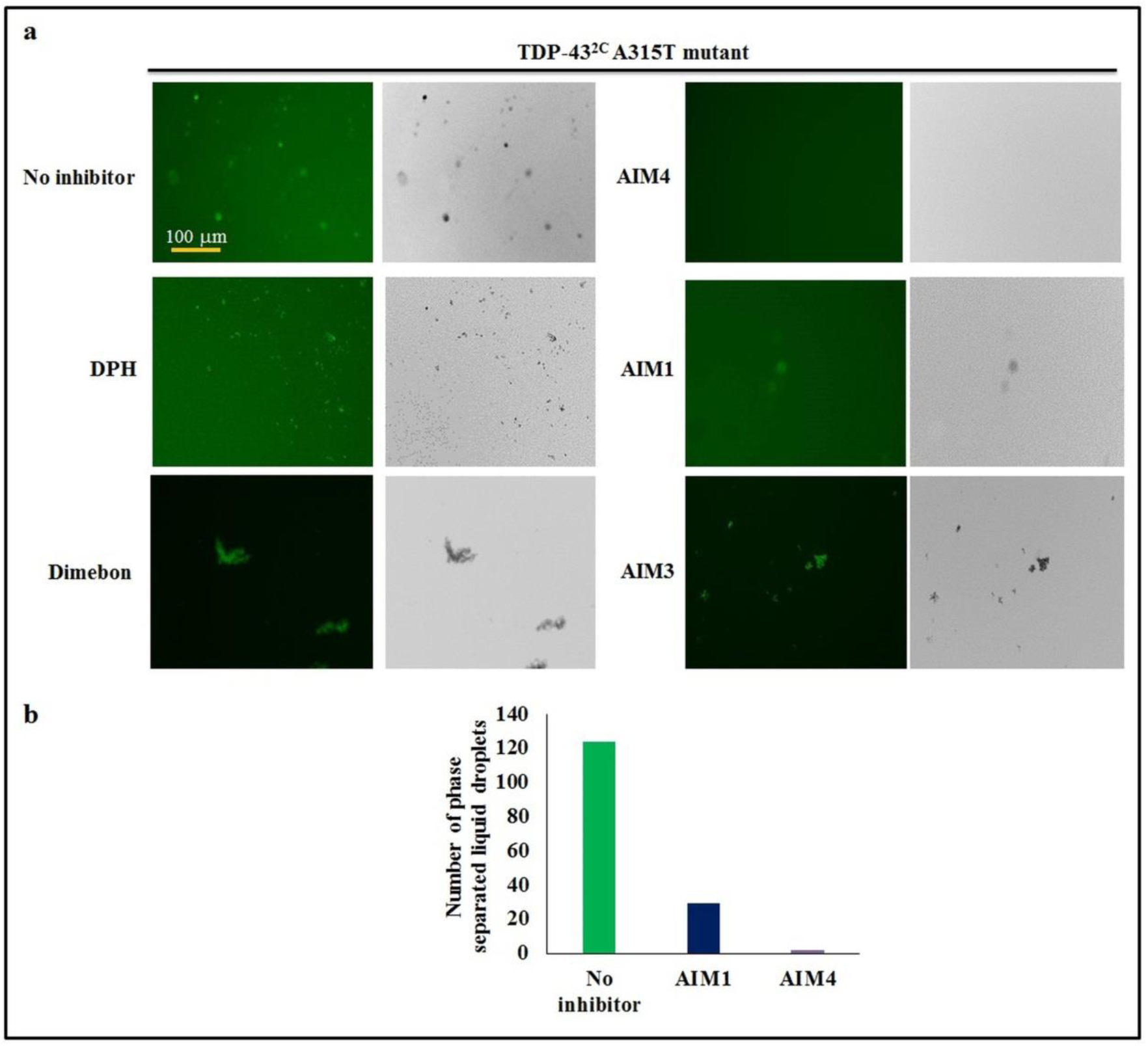
Acridine derivative AIM4 inhibits the *in vitro* liquid-liquid phase separation (LLPS) of the TDP-43^2C^-A315T protein. **a**. Mutant A315T TDP-43^2C^, labelled with Alexa Fluor™ 488 C5 maleimide undergoes transition from soluble state to liquid droplets suggesting of a process termed as liquid-liquid phase separation (LLPS). Mutant TDP-43^2C^-A315T (400 µM) was incubated at 37 °C in aggregation buffer (2.5M urea-PBS), either alone or with the additions of 1:15 (protein: compound molar ratio) of AIM4, AIM3, AIM1, DPH or dimebon, for a period of 5 hours. The Alexa Fluor-bound species appear as droplets as observed under fluorescence microscopy when examined using the GFP filter. With acridine compounds, AIM3 which affected TDP-43^2C^ aggregation to the least extent exhibited the formation of irregular solid shaped aggregates, whereas AIM1 which moderately inhibits TDP-43^2C^ aggregation, the formation of these liquid droplets gradually decreased and were present in far fewer numbers in comparison with the protein sample without any inhibitor and with AIM4, which exhibits maximum inhibitory potential to TDP-43^2C^, the liquid droplets completely disappeared thereby indicating a complete inhibition of LLPS. All the images were taken at 10X magnification. **b**. The total number of liquid droplets found in each frame of the Alexa Fluor labelled protein without any inhibitor as well as with 1:15 (protein: compound molar ratio) of AIM1 and AIM4 were counted. The data represents the number of droplets found in 18 frames for each of the sample.

### Conclusion

Owing to the importance of inhibition of TDP-43 aggregation and the current unavailability of effective drugs against ALS, a small molecule AIM4 had been previously demonstrated to be a candidate for the inhibition of the TDP-43 aggregation [71]. Here, mechanism of AIM4-mediated inhibition was investigated through computational tools and it was further examined if AIM4 can also inhibit the mutant TDP-43^2C^-A315T’s amyloid-like aggregation and liquid-liquid phase separation, the potential precursor process leading to its aggregation [99]. Recently, a cryo-EM structure was solved from the core amyloidogenic region (aa: 288-319; PDB ID: 6N3C) of TDP-43 having A315E familial ALS mutation [75] and here another familial A315T mutation was incorporated *in silico* into the 6N3C structure and these two structures were used for examining AIM4’s affinity. In addition, other known structures from TDP-43 sequence namely, tandem RRMs (aa: 96-269; PDB ID: 4BS2), N-terminal domain (aa: 1-89; PDB ID: 2N4P) and ten amyloidogenic peptide fragments from the low complexity domain (LCD) of TDP-43, were also used to predict the binding site of AIM4. Computational studies predicted better binding of AIM4 than the other used ligands to the aa: 288-319 amyloidogenic core region of TDP-43 with the A315E and A315T familial mutations. Also, from the *in silico* studies, in the 6N3C-A315T mutant TDP-43 structure (aa: 288-319), AIM4 was predicted to interact preferentially with two amino acids, glycine and phenylalanine which are also already known to be involved in its pathogenic liquid-liquid phase separation (LLPS). Therefore, the ability of AIM4 to modulate LLPS of TDP-43 was investigated *in vitro* using a recombinantly purified C-terminal fragment, TDP-43^2C^-A315T containing aa: 193-414 of TDP-43. Strikingly, AIM4 was found here to inhibit the LLPS of the TDP-43^2C^-A315T protein. In summary, this study predicted the C-terminal region aa: 288-319 of TDP-43 to have a possible binding site for AIM4 which may be responsible for the inhibition of TDP-43 aggregation by AIM4. Furthermore, of high significance, the study also demonstrated an ability of AIM4 to inhibit the *in vitro* LLPS of the mutant A315T-TDP-43^2C^ proteins thereby asserting the need for further investigations on applicability of AIM4 as an anti-TDP-43 aggregation molecule with relevance to ALS therapeutics.

## Acknowledgements

We thank IIT Hyderabad funded by MHRD, Govt. of India for research infrastructure and support. AG and GR are thankful to MHRD, Govt. of India, for senior research fellowship (SRF). VB thanks DBT, Govt. of India, for SRF. VRT thanks MHRD for research assistantship. SA and USM thank MHRD for junior research fellowship (JRF). Basant K Patel thanks DST, Govt. of India for research grant (Grant no: EMR/2016/006327).

