## Supplementary figures for "Computational insights into mechanism of AIM4-mediated inhibition of aggregation of TDP-43 protein implicated in ALS and evidence for *in vitro* inhibition of liquid-liquid phase separation (LLPS) of TDP-43^2C^-A315T by AIM4"

**Title**

**Affiliation**

### Supplementary Figures

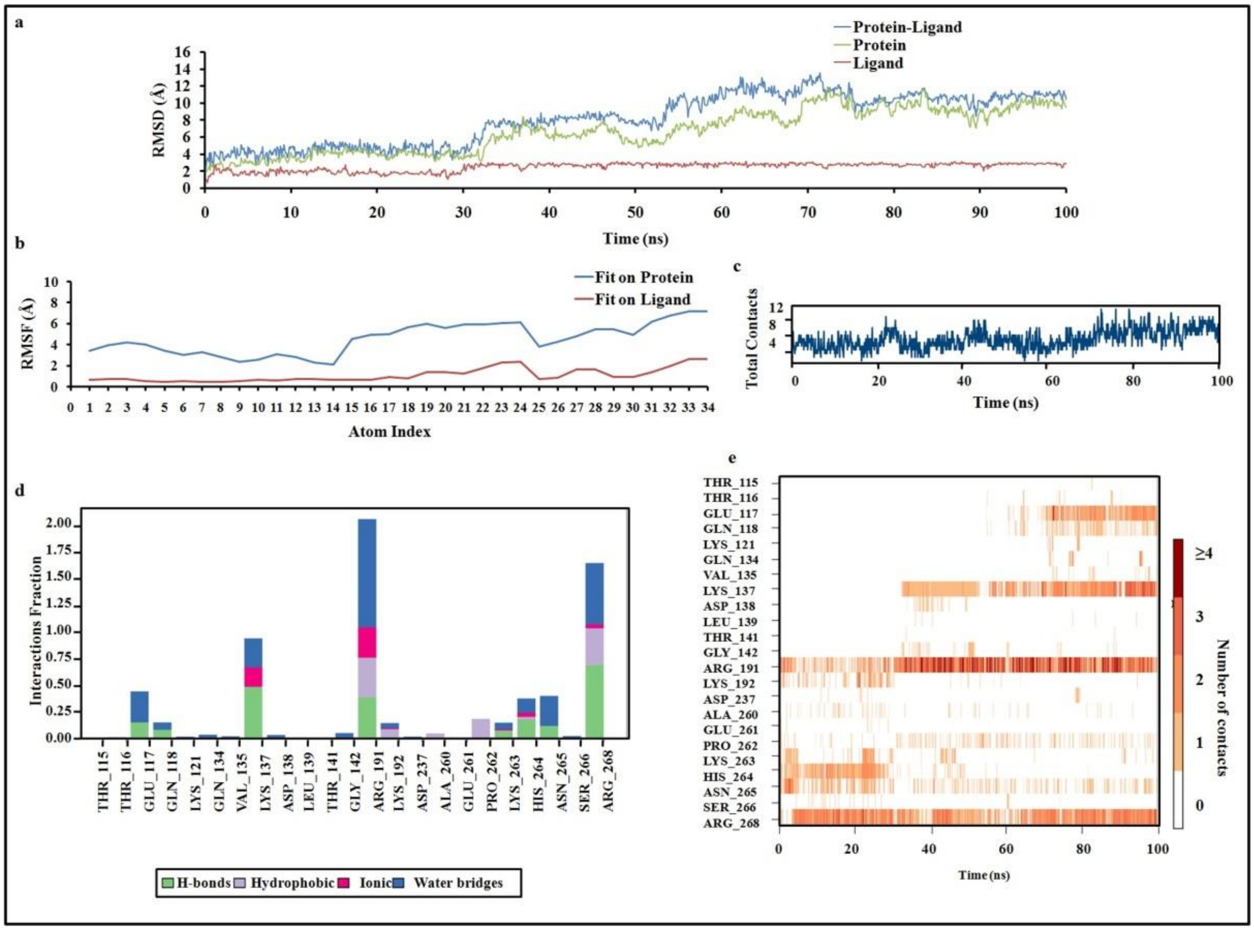


**Figure S1: MD simulation (100ns) of the complex of AIM4 with the structure of the TDP-43's tandem RRMs (aa: 96-269; PDB ID: 4BS2).**

**a.** RMSD of the protein, ligand and protein-ligand complex during MD simulation. For the 4BS2-AIM4 complex, the RMSD was not found to be plateaued till the end of the simulation indicating that the complex is not stable. **b.** Root Mean Square Fluctuations (RMSF). 'Fit Ligand on Protein' trend indicates the ligand fluctuations with respect to the protein. 'Ligand' trend shows the fluctuations where the ligand in each frame is aligned on the ligand in the first reference frame **c.** The number of total contacts (H-bonds, Hydrophobic, Ionic, Water bridges) made between the TDP-43-RRMs and the ligand (AIM4) over the course of the simulation. **d.** The interactions of AIM4 with the amino acids of TDP-43-RRM1-2 (aa: 96-269; PDB ID: 4BS2). The stacked bar charts have been normalized over the course of the trajectory: for example, a value of 1.0 suggests that the specific interaction is maintained for 100% of the simulation time. **e.** A timeline representation of the interactions and contacts (H-bonds, Hydrophobic, Ionic, Water bridges). it shows the residues of TDP-43-RRM1-2 interacting with the ligand (AIM4) in each trajectory frame. Some residues make more than one specific contact with the ligand, which is represented by a darker shade of orange, according to the scale to the right of the plot.

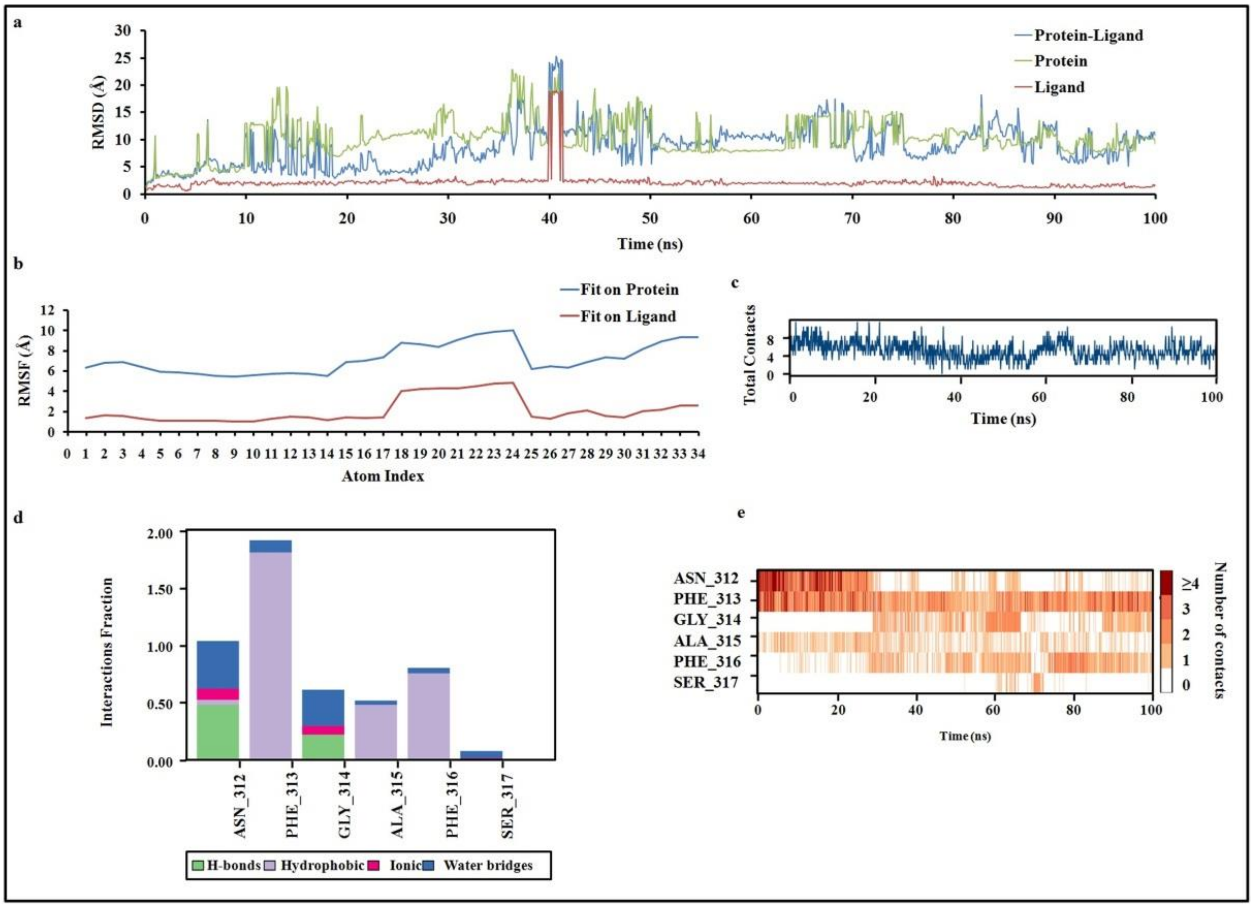


**Figure S2: MD simulation (100ns) of the complex of AIM4 with short amyloidogenic peptide from LCD of TDP-43 (aa: 312-317; PDB ID: 5WHN).**

**a.** RMSD of the protein, ligand and protein-ligand complex during MD simulation. For the 5WHN-AIM4 complex, significant high fluctuations were observed in the protein and protein ligand RMSD. Also the RMSD was not found to be plateaued till the end of the simulation indicating that the complex is not stable. **b.** Root Mean Square Fluctuations (RMSF). 'Fit Ligand on Protein' trend indicates the ligand fluctuations with respect to the protein. 'Ligand' trend shows the fluctuations where the ligand in each frame is aligned on the ligand in the first reference frame **c.** The number of total contacts (H-bonds, Hydrophobic, Ionic, Water bridges) made between the 5WHN peptide and the ligand (AIM4) over the course of the simulation. **d.** The interactions of AIM4 with the amino acids of 5WHN peptide (aa: 312-317). The stacked bar charts have been normalized over the course of the trajectory: for example, a value of 1.0 suggests that the specific interaction is maintained for 100% of the simulation time. **e.** A timeline representation of the interactions and contacts (H-bonds, Hydrophobic, Ionic, Water bridges). it shows the residues of 5WHN peptide interacting with the ligand (AIM4) in each trajectory frame. Some residues make more than one specific contact with the ligand, which is represented by a darker shade of orange, according to the scale to the right of the plot.

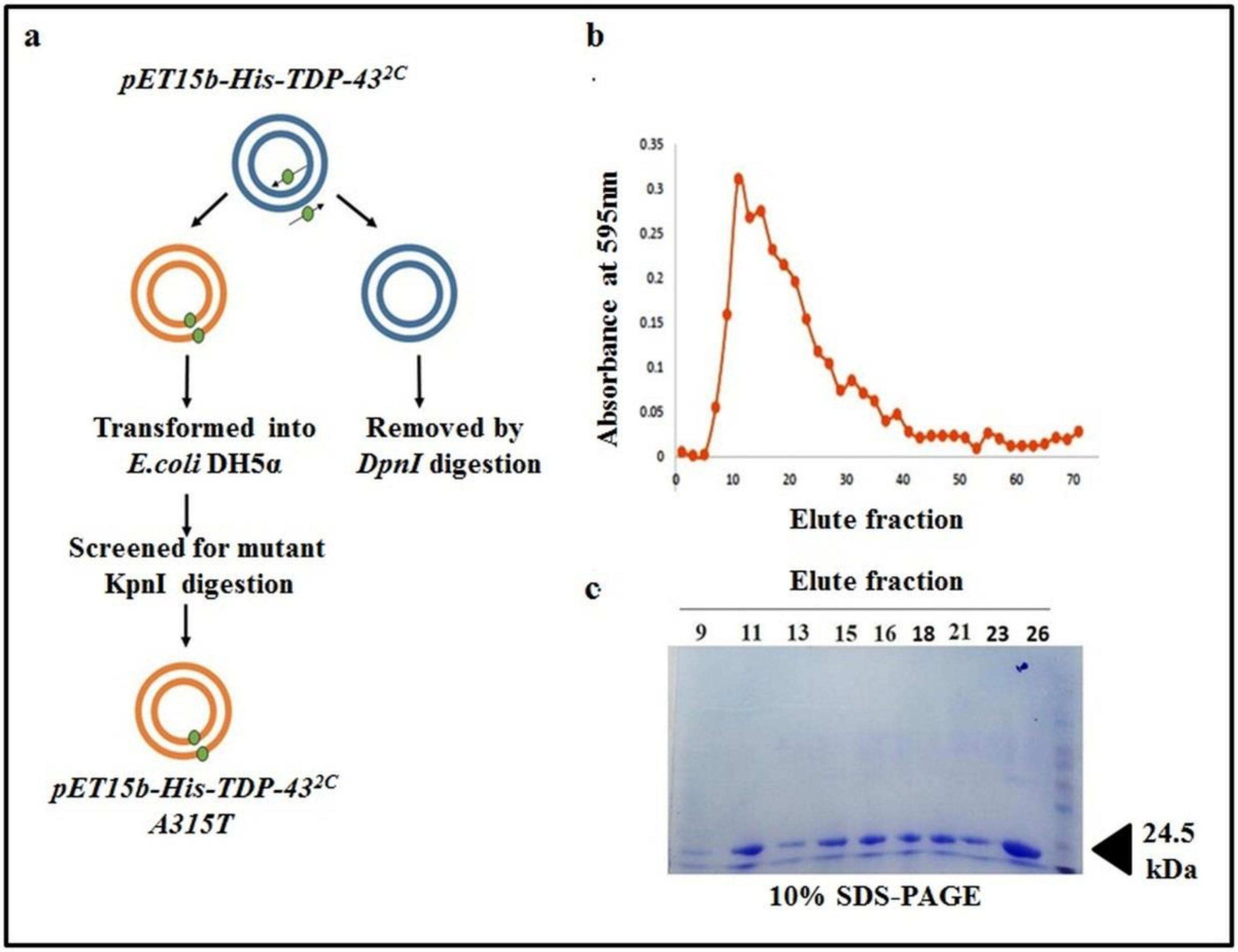


**Figure S3:** ***In vitro* mutagenesis and recombinant mutant TDP-43$^{2C}$-A315T protein purification.**

**a**. Primers were designed to incorporate A315T mutation in TDP-43$^{2C}$ using plasmid *pET15b-His-TDP-43$^{2C}$* as template and the parental strand was removed by digestion with *Dpn1*. After digestion, plasmid was transformed into *E.coli* DH5α and positive mutagenic clone was confirmed by *Kpn1* digestion. *pET15b-His-TDP-43$^{2C}$ A315T* plasmid was transformed into Rosetta cells for Ni-NTA protein purification and the concentration and purity of purified protein in elute fraction was measured by using **b.** Bradford assay and **c.** SDS-PAGE analysis, respectively.
